# The functional organisation of the hippocampus along its long axis is gradual and predicts recollection

**DOI:** 10.1101/428292

**Authors:** Izabela Przezdzik, Myrthe Faber, Guillén Fernández, Christian F. Beckmann, Koen V. Haak

## Abstract

Understanding the functional organisation of the hippocampus is crucial for understanding its role in cognition and disorders in which it is implicated. Different views have been proposed of how function is distributed along its long axis: one view suggests segregation, whereas the alternative view postulates a more gradual organisation. Here, we applied a novel ‘connectopic mapping’ data-analysis approach to the resting-state fMRI data of participants of the Human Connectome Project, and demonstrate that the functional organisation of the hippocampal longitudinal axis is gradual rather than segregated into parcels. In addition, we show that inter-individual variations in this gradual organisation predicts variations in recollection memory better than a characterisation based on parcellation. These results present an important step forward in understanding the functional organisation of the human hippocampus and have important implications for translating between rodent and human research.

## Introduction

The hippocampus is involved in multiple cognitive functions including episodic memory (Scoville and Milner, 1957; Squire, 1992), spatial navigation (Morris et al., 1982; Maguire et al., 1998), and emotion-related processing (Gray and McNaughton, 2000; Bannerman et al., 2004). Despite decades of research, it is still unclear how its macroscopic organisation subserves these multiple cognitive functions. Although there is consensus that the hippocampus is functionally organised along its longitudinal axis, different views have been proposed of how function is distributed along this axis: one view suggests that the neural circuits associated with different functions are segregated into discrete hippocampal subdivisions with sharply demarcated borders, whereas the alternative view postulates a gradual organisation of function along the long axis (Strange et al., 2014). Distinguishing between these alternative views is important, because these two alternative characterisations of the underlying neurobiology may lead to very different approaches when analysing signals recorded from the hippocampus and will certainly lead to different interpretations of hippocampal function, as we will demonstrate in this paper.

Early anatomical studies (Swanson and Cowan, 1977), electrophysiological (Elul, 1964; Racine et al., 1977) and lesion studies in rodents (Henke, 1990; Moser et al., 1993) found differences in cortical and subcortical projections from the dorsal and ventral hippocampus, lending support to the idea that the hippocampus can be parcellated into functionally-distinct subdivisions (for an extensive review, see Strange et al., 2014). Consequently, multiple proposals attempting to allocate alternative functions to the ventral and dorsal portions—which correspond to anterior and posterior sections of the hippocampus in humans—have been introduced (see e.g. Poppenk et al. 2013), suggesting that the ventral (anterior) portion is primarily involved in emotion-related processing and the dorsal (posterior) in memory and spatial processing (Strange et al., 2014). However, recent tracing studies have shown that the cortical input from the entorhinal cortex to the hippocampus can be described in terms of a dorsal - ventral gradient (Witter et al., 2000). In addition, hippocampal place cells can be found along the entire extent of the longitudinal axis of the hippocampus, with their field size increasing gradually from the dorsal to ventral sub-regions, demonstrating a scale-related gradient of functional change within the hippocampus (Kjelstrup et al., 2008).

It is important to note that the parcellated- and gradient views are not necessarily mutually exclusive: it is possible that multiple functional gradients are superimposed on discrete hippocampal functional domains (Strange et al., 2014). Studies that could shed light on this are markedly lacking in the field, in particular in human neuroscience. One of the main reasons why the gradient-like organisation of the hippocampus has been under-explored in humans is a lack of appropriate methods: the invasive nature of tracing studies that have first suggested a gradient render them unsuitable for human participants. Studies into the functional organisation of the human hippocampus have therefore predominantly been based on parcellation-based approaches that rely on non-invasive brain imaging techniques. However, by using parcellation methods, one forces the characterisation of functional organisation to be in terms of strictly segregated parcels, even if the true functional organisation is smooth without sharp borders.

Therefore, we here set out to investigate the functional organisation of the human hippocampus using ‘connectopic mapping’, an emergent approach to characterising functional organisation non-invasively in individual human participants without imposing a parcellation scheme (Haak et al., 2017). Connectopic mapping specifically aims at characterising gradual changes in the location-dependent pattern of associated functional connectivity. Here, we test if the functional organisation of the human hippocampus in terms of the location-depend pattern of functional connectivity might be more meaningfully described as a gradient than in terms of parcels. We do that by testing whether inter-individual variations in the gradient predict inter-individual variations in hippocampus-related behaviour better than a parcellation-based approach.

## Materials and Methods

### rfMRI data and pre-processing

A data-set comprising participants of the WU-Minn Human Connectome Project (S-500 release) (Van Essen et al., 2013) was used in this study. In the connectopic mapping analysis we included only those participants (N = 475) who completed all four resting-state fMRI (rfMRI) sessions (multi-band, TR = 0.72s). The within-run resting-state data were pre-processed as detailed in Smith et al., 2013 including spatial distortions and head motion correction, T1w registration, resampling to 2 mm MNI space, global intensity normalisation, high-pass filtering with a cut-off at 2000s, and the ICA-based artefact removal procedure (FSL-FIX, Griffanti et al., 2014; Salimi-Khorshidi et al., 2014). In addition, before applying connectopic mapping we smoothed the data with a 6 FWHM Gaussian kernel, regressed the mean ventricular as well as white-matter signal from the time-series, and normalise it. Finally, we concatenated the data from four resting-state scans into 1-hour session. These pre-processed data were then used to estimate connectopic maps for each individual (Haak et al., 2017).

### Connectopic mapping

Connectopic mapping (Haak et al., 2017) is a data-driven approach for mapping the connectopic organisation of brain areas based on resting-state functional magnetic resonance imaging (rfMRI) data. Previous work has shown that this method accurately traces known functional gradients in brain regions such as retinotopic and somatotopic cortex, as well as striatum and entorhinal cortex (Haak et al., 2017; Navarro Schröder et al., 2015, Marquand et al., 2017) Furthermore, recent work has linked inter-individual differences in cortico-striatal connectopic organisation to meaningful variations in goal-directed behaviour (Marquand et al., 2017).

Details of the connectopic mapping procedure are described in Haak et al. 2017. Briefly, for every voxel in the region of interest (ROI; here, the left or right hippocampus), we obtained a “connectivity fingerprint” by computing the correlation between the voxel-wise time-series and the rest of the cortex (a singular value decomposed matrix of time-series of all grey-matter voxels outside the ROI). We then computed the within-ROI similarity of functional connectivity, and applied non-linear manifold learning (Laplacian Eigenmaps) to the graph representation of this similarity matrix to obtain the connectopic maps, indicating how hippocampal-neocortical connections vary topographically across the ROI.

Connectopic mapping was applied to the resting-state fMRI data of 475 participants. Hippocampal ROIs (one for each cerebral hemisphere) were based on the Harvard-Oxford atlas. As a result, we obtained the connectopic maps describing each participant’s hippocampal-neocortical functional connectivity patterns for left and right hippocampus separately. The connectopic maps of interest were captured by the eigenmaps associated with second-smallest eigenvalue, which were then used in all subsequent analyses.

### Trend surface modelling

In order to enable statistical inference over the connectopic maps we used trend surface modelling (Haak et al. 2017). This involves fitting series of polynomial basis functions to the connectopic maps to capture their overall spatial pattern in a small number of coefficients. A spatial model of the dominant connectopic map was estimated for each participant and hemisphere independently.

We started the estimation with fitting a polynomial of degree 1 (a straight line with a slope) and investigated progressively more refined approximations, by combining the lower order models up to the fifth model order. Because hippocampi are three-dimensional structures, this entails estimation within the x, y and z direction, resulting in three trend surface model parameters (TSM parameters) capturing the gradient’s overall spatial pattern in the first model order estimation. The second model order entails estimation in the same directions but fitting the polynomial of degree 2 (a <u>parabola)</u>. After combining this with the estimates of lower polynomial basis functions, it results in six parameters that refer to x, y, z, x^2^, y^2^, z^2^. Accordingly, the number of parameters increases as we move to the higher order of the trend surface models.

We fitted these models using the Bayesian linear regression, which also yielded estimates of the likelihood of the model given the data, based on which we later computed the Bayesian Information Criterion (BIC) and Akaike Information Criterion (AIC) scores for model order selection purposes.

### Behavioural data

For testing associations between inter-individual differences in connectivity gradients and subject-dependent behaviour we derive a surrogate measure of recollection performance. HCP participants performed a series of tasks during separate fMRI scanning sessions, including an N-back task in which four different stimulus types (pictures of faces, places, tools and body parts) were shown in separate blocks. After completing the *N*-back task in the scanning session, each participant’s memory was tested using a Remember-Know paradigm (Tulving, 1983; 1985). Participants were presented with the images of faces and places earlier presented in the N-back task, mixed with an equal number of foil items. The body parts and tools were not included in the testing set, as there were not enough new items to create foil stimuli for those categories (see Barch et al. 2013). For each item, participants reported whether they had seen it before (old-new discrimination), and for each item that was reported as old, they were asked to indicate whether they could recollect the encoding context of the item (“Remember”-response) or not (“Know”-response). The “Remember” and “Know” responses are thought to reflect different, independent processes as evidenced by neuroimaging research that has shown that “Remember” responses are hippocampus - dependent, whereas “Know” responses rely on higher-order visual processing areas (Eldridge et al., 2000).

We computed *d*-prime (*d*’) measures of old-new discrimination (recognition), and excluded participants whose *d*’ was at or below zero (i.e., participants with below chance performance in either the face or place condition or both) from further analysis (16 subjects were excluded based on below-chance performance in the face condition, 4 for below-chance performance in the place condition). Three additional participants with missing behavioural data were also excluded from further analysis. This resulted in *N* = 448 (265 females; 22–36 years, mean age = 29.21, SD = 3.50) subjects for analysis of the face condition, and *N* =460 (271 females; age, 22–36 years, mean age = 29.16 years; *SD* = 3.51 years) subjects for analysis of the place condition. To isolate hippocampus-mediated recollection from more generic recognition (as measured by *d*’), we computed the inverse of the independence remember/know equation (Jacoby et al., 1997): Recollection = proportion of “Remember” responses / 1-proportion of “Know” responses. In its original form, this formula quantifies the contribution of familiarity-based recognition (i.e., recognising an item but not recollecting its encoding context) to overall memory performance. The inverse represents the proportion of recollection over and above recognition, and therefore specifically taps into the hippocampal mechanisms that underlie retrieval of episodic detail. This measure was used as the dependent variable in subsequent analyses.

## Statistical analysis

### General linear model (GLM)

A GLM approach was used to investigate whether the TSM parameters, which quantitatively describe the hippocampal connectivity gradients derived from resting-state fMRI at an individual level, predict recollection memory. The recollection scores for faces and places were used as dependent variables in two separate models. Age, head movement during the scan (mean frame-wise displacement), and the reconstruction algorithm version that was used for reconstruction the rfMRI data from k-space were added as covariates (the latter changed during HCP data collection and has a substantial influence on rfMRI connectivity estimates). As we were interested in the variance explained by the TSM parameters *over and above* the variance explained by the covariate variables, we computed the partial *R*^2^ as (*RSS*_reduced_ – *RSS*_full_) / *RSS*_reduced_. Accordingly, in the full model we included the TSM parameters, age, motion, and the reconstruction method, whereas the reduced model included only age, motion, and the reconstruction method. The same approach was used to test if the TSM models predict *d*’ measures of old-new discrimination. A permutation testing procedure implemented in FSL-PALM that accounts for the family structure of the HCP sample (Winkler et al., 2015) was used to assess the statistical significance of the ensuing partial *R*^2^ values (with 5K permutations).

### K-means clustering

Parcellation approaches have suggested a positive relationship between recollection and posterior hippocampus volume, in particular when expressed as a ratio to anterior hippocampus volume (e.g., Poppenk & Moscovitch, 2011). We therefore tested whether the gradient-organisation of hippocampus explains individual differences in recollection over and above parcellation. We used k-means clustering to obtain anterior and posterior parcels, and computed the ratio between them, approximating previous parcellation studies (Poppenk and Moscovitch 2011). We used the ratio as a predictor in the GLM analysis to test whether it (over and above the covariates: age, motion and reconstruction version) can on its own predict recollection memory. We then tested whether the TSM parameters explain variance over and above this model. Lastly, we tested whether both the ratios and TSM estimates of the gradients, can explain substantially more variance in the recollection score than the TSM estimates alone.

## Results

### Gradual functional connectivity patterns within the human hippocampus

Connectopic mapping was applied to the resting-state fMRI data to estimate the hippocampal-neocortical functional connectivity patterns in the hippocampus at the individual level. At the group-level, the dominant connectopic map, which represents the first dominant mode of the connectivity change, followed the expected anterior-to-posterior trajectory (Figure 1).

**Figure 1.**
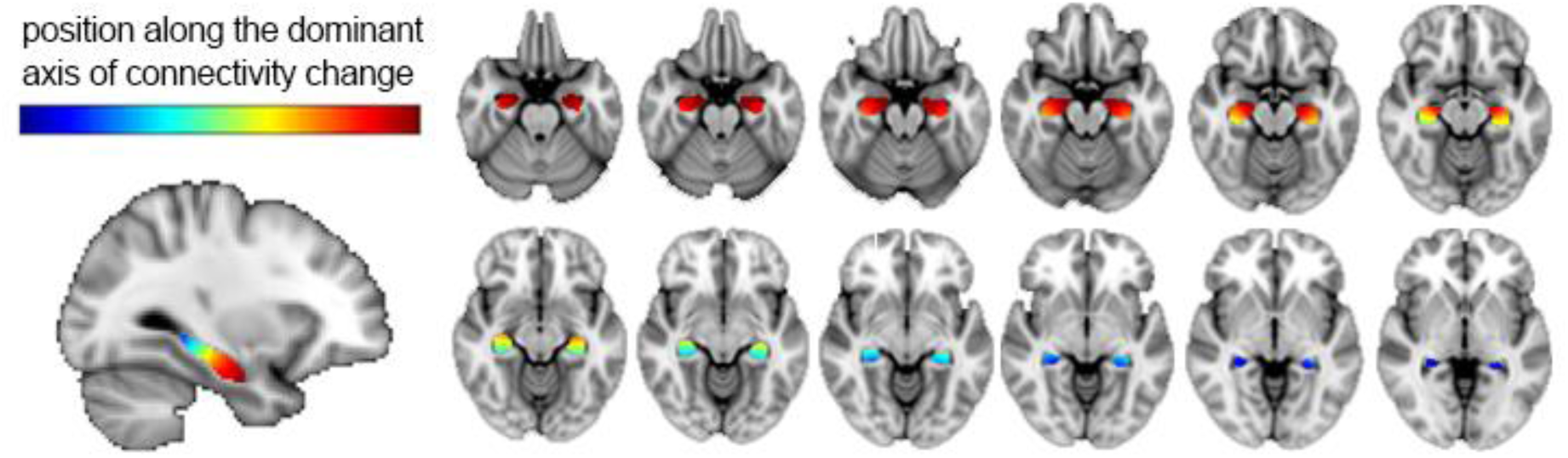
The hippocampal-neocortical connectivity gradient at the group level (N=475) stretches along the long hippocampus axis. The colour bar indicates the position along the dominant mode of connectivity change, and so similar colours represent similar connectivity patterns. Changes in colour represent changes in topographically organised functional connectivity. The values within this arbitrary range, as a gradient represent connectivity change.

Figure 2 shows each individual’s dominant connectopic map as a function of the Euclidian cortical distance from the hippocampus’ most posterior voxel (for the left and right hippocampus separately). Notably, the connectopic changes are rather smooth and gradual, without any fast transitions. Thus, the connectopic mapping results indicate a gradual change along the long hippocampus axis that follows an anterior-to-posterior trajectory, resembling previously reported findings from animal studies that showed a ventral-dorsal gradient-like organisation in rodent hippocampus. Figure 2 further indicates that while the overall pattern across the participants looks very similar, the patterns are not identical between the participants. The spatial properties of these single-level gradients were further estimated with the trend surface modelling and used to predict recollection.

**Figure 2.**
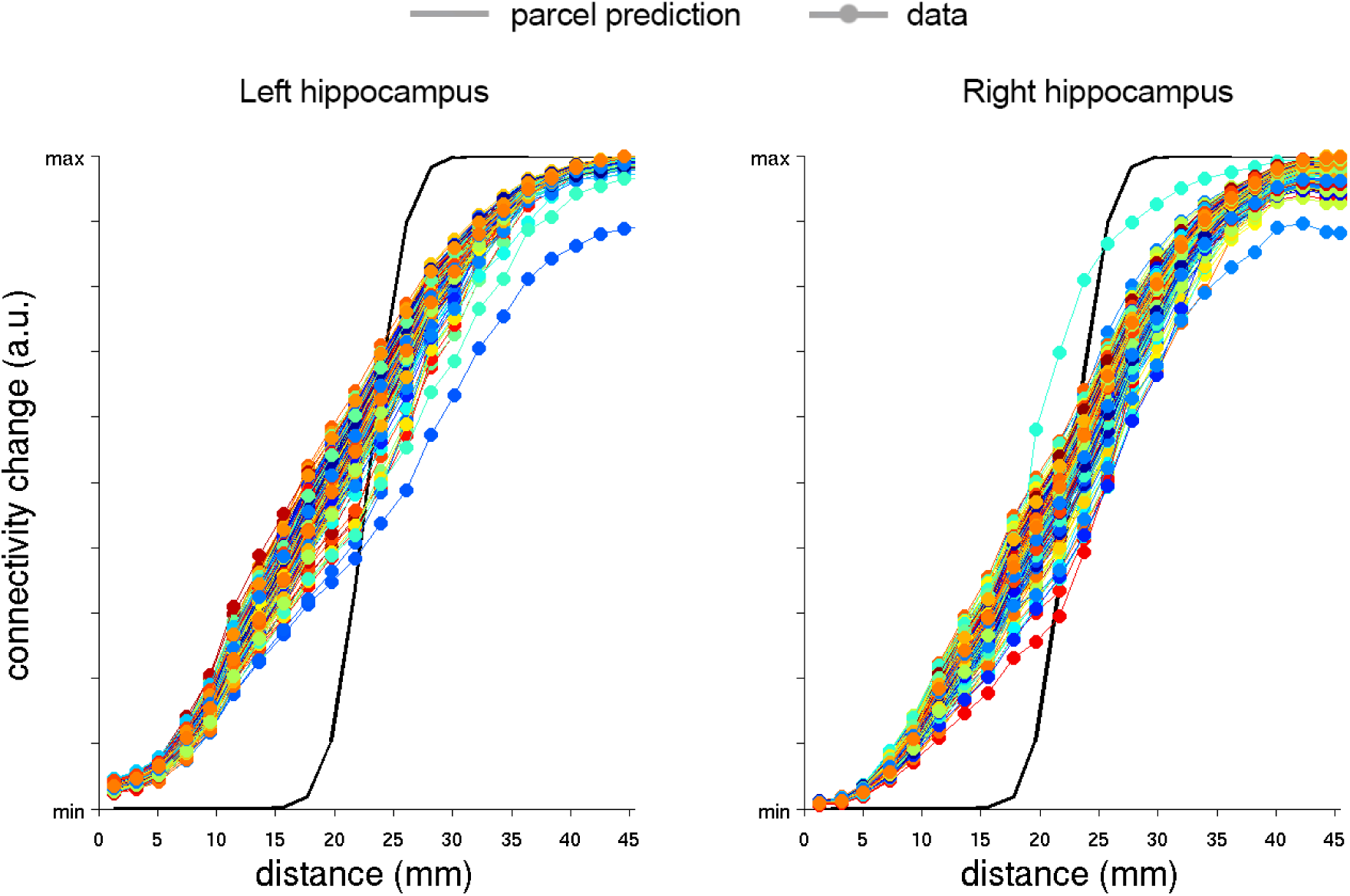
Left and right hippocampal-neocortical functional connectivity patterns plotted against the distance from the most posterior voxel in the hippocampus. Data were binned in terms of distance (23 bins of ~2mm). Each coloured line represents one participant. The black line represents a non-gradient where the transition is fully induced by smoothing discrete parcels using the same Gaussian kernel.

### Feature extraction: Trend surface modelling analysis of the connectopic maps

In order to investigate whether individual differences in the obtained connectivity gradients are functionally meaningful, we first reduced the number of estimates characterising the connectopic maps by employing trend surface modelling (TSM), which summarises the overall voxel-wise spatial pattern of the individual connectopic maps in a small number of spatial model parameters. From the series of trend surface models that were fitted, the Bayesian Information Criterion (BIC) indicated that the third and fourth TSM model orders were most favourable, with only very little difference between them in terms of variance explained (average across hemispheres 98.65% and 98.75%, respectively; see Fig. 3; the Akaike Information Criterion (AIC) showed similar results). We therefore report the results for both model orders.

**Figure 3.**
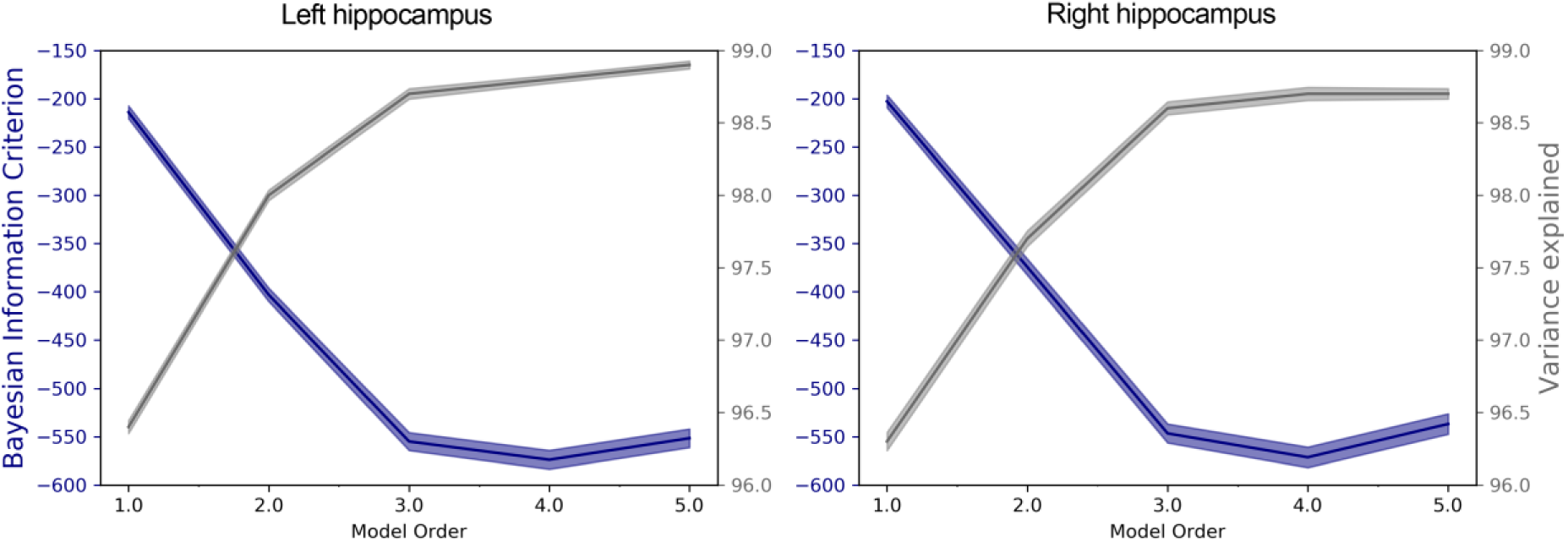
TSM model selection. The average values of the Bayesian Information Criterion (BIC) and the proportion of the variance of the overall spatial pattern explained by the respective trend surface model orders. The shaded area represents the 95% confidence intervals.

## Behavioural results

The average of *d*’ scores (including scores of those participants that had *d*’ scores equal or less than zero, N = 464) collapsed across both stimulus types was 1.37 (SD = 1.24), indicating that on average the participants performed the task well. The average *d*’ for faces was 1.06 (SD = 0.97) and the average *d*’ for places was 1.69 (SD = 1.50). The difference between *d*’ faces and *d*’ places was statistically significant (average difference = 0.63, p = 0.00), where statistical significance was assessed using FSL**-**PALM (5K sign-flipping). The hippocampus-mediated recollection scores were calculated based on the inverse of the independence remember/know (IRK) equation (see Methods), which ranges between 0 and 1. The average recollection score for faces was 0.61 (SD = 0.22), whereas the average recollection score for places was 0.46 (SD = 0.22). This difference was statistically significant (average difference = 0.14, p = 0.00), generated by FSL-PALM (5K sign-flipping), indicating that faces were recollected more often than places. Since there are differences between these behavioural measures in terms of old-new discrimination and recollection, we treat them separately in subsequent analyses.

### Recollection memory in relation to connectopic mapping

We used a GLM to investigate whether TSM parameters that summarise the gradients at the individual level predict hippocampal-dependent recollection. We found that indeed, recollection was significantly predicted by TSM parameters (3^rd^ order, nine parameters) over and above the covariates, such that the left hippocampal connectivity gradient predicted recollection memory for faces (Partial R^2^ = 0.057, p = 0.002, grey bar in Figure 4A, below the exemplary image of the stimulus type: faces), and the right hippocampal connectivity gradient predicted recollection for places (Partial R^2^ = 0.041, p = 0.032, grey bar in Figure 4A, below the exemplary image of the stimulus type: places). A similar pattern was found when the gradient was approximated with a 4^th^ model order and 12 parameters (these results are not presented in the figure): the left hippocampal connectivity gradient was significantly predictive of recollection for faces (Partial R^2^ = 0.063, p = 0.006), whereas the right hippocampal connectivity gradient showed a relationship with recollection for places, albeit marginally significant (Partial R^2^ = 0.042, p = 0.092). These results suggest that the gradient-like functional organisation of the hippocampus at the individual level is predictive of individual differences in recollection.

**Figure 4.**
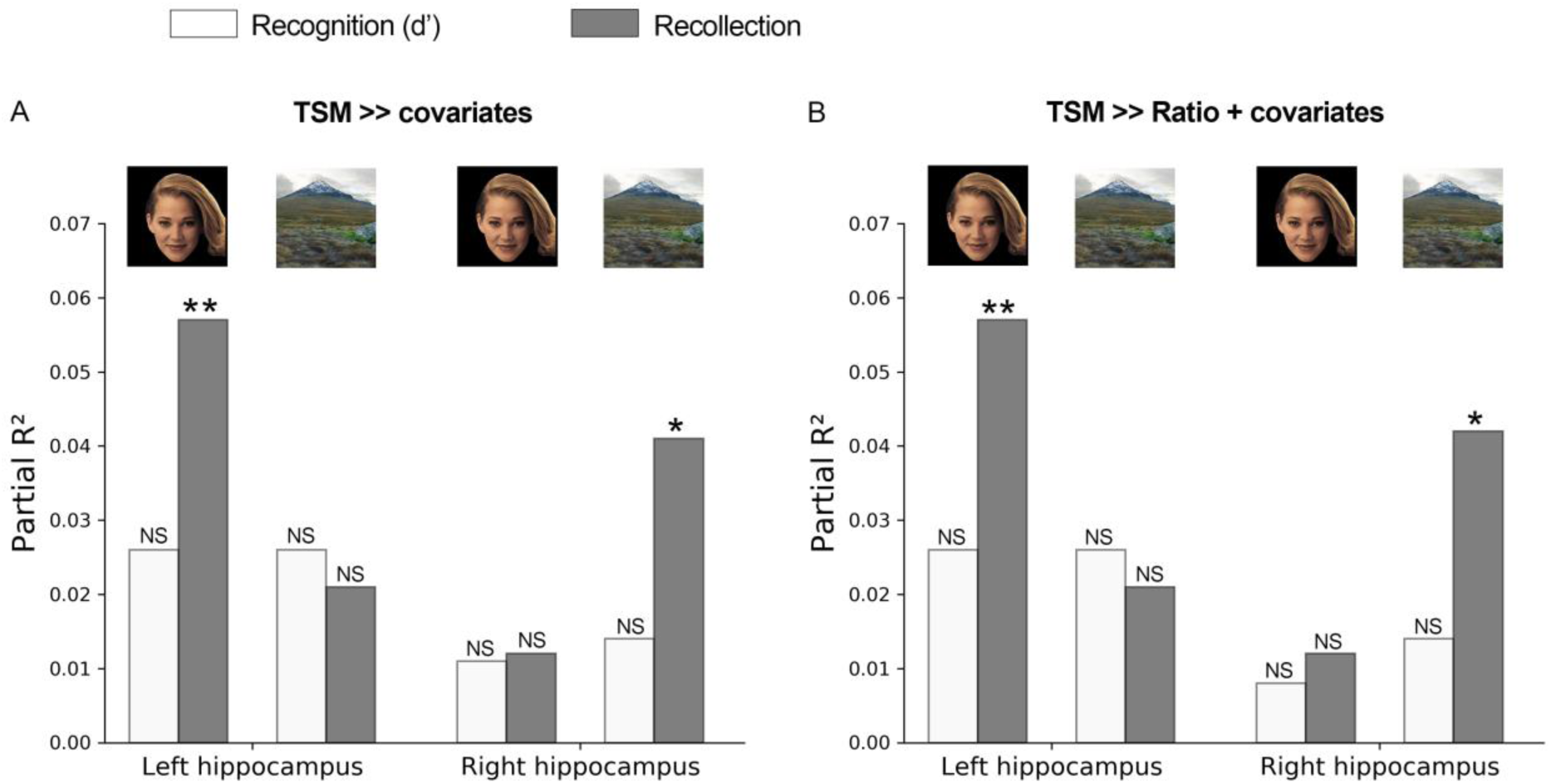
Proportion of the variance of the recognition (*d*’) and recollection scores explained by the spatial model coefficients (TSM) over and above the variance explained by age, head motion and fMRI reconstruction algorithm version. B. Proportion of the variance of the recognition (*d*’) scores and recollection scores explained by the spatial model coefficients (TSM) over and above the variance explained by age, head motion, fMRI reconstruction algorithm version and the parcel’s ratio. Model order refers to the model order of the trend surface model that was fitted to each individual’s hippocampal connectopic map. Here, the results are presented for the third model order. * p < 0.05, ** p< 0.01.

As a sanity check, we investigated whether TMS parameters are associated with old-new discrimination (*d*’ faces and *d*’ places). As predicted, the GLM results for both third and fourth model order showed that the TSM parameters of the gradient in neither the left nor right hippocampus predicted recognition (discrimination between old-new items) for places and faces (all p ≥ 0.204); white bars in Figure 4A).

### Parcels vs. gradient along the long axis

Previous studies showed a relationship between recollection and the ratio of posterior vs anterior hippocampus (e.g., Poppenk & Moscovitch, 2011). It is possible that the gradient explains the same variance in recollection as these ratios. In that case, TSM parameters should not explain variance over and above the posterior vs. anterior ratios. It is also possible that the TSM parameters and ratios each explain unique variance, which means that a model that contains both the ratios and TSM parameters explains most variance, suggesting a superposition of a gradient on top of a parcellation. However, if the TSM parameters — and critically, not the ratios — explain individual differences in recollection (i.e., a significant increase in variance explained when adding the TSM parameters to the ratio model but not vice versa), the findings would suggest that a description of the functional organisation of the hippocampus in terms of a gradient is more functionally meaningful than a description in terms of parcels.

We therefore first tested whether the ratios, obtained by splitting the functional connectivity gradient into two parcels and computing the ratio of the posterior vs anterior part, predicted memory performance. We found that neither recognition (*d*’) scores nor recollection scores displayed a significant relationship with the posterior vs. anterior ratios (all p ≥ 0.19). Adding the TSM parameters to the ratio model explained a significant amount of variance over and above the ratios. The results are summarised in Figure 4B (3rd model order: left hippocampal connectivity gradient predicted recollection for faces (Partial R^2^ = 0.057, p = 0.003; grey bar below the exemplary image of the stimulus type: faces), and the right hippocampal connectivity gradient predicted recollection for places (Partial R^2^ = 0.042, p = 0.027, grey bar below the exemplary image of the stimulus type: places; 4^th^ model order: the left hippocampal connectivity gradient and recollection for faces (Partial R^2^ = 0.063, p = 0.005), the right hippocampal connectivity gradient and recollection for places, (Partial R^2^ = 0.042, p = 0.094). Vice versa, adding the ratios to a model that predicts recollection from the TSM parameters does not significantly increase the explained variance (all p ≥ 0.18), suggesting that inter-individual differences in the gradient, rather than inter-individual differences in the posterior-anterior ratio, are related to recollection.

## Discussion

In the present study, we tested whether inter-individual differences in the gradual change of topographically organised, hippocampal-neocortical functional connectivity predicts hippocampus-dependent recollection, over and above a parcellated view. We used a novel data analysis approach, connectopic mapping, which revealed a smooth connectivity gradient that follows the anterior to posterior trajectory of the longitudinal hippocampus axis. After estimating these gradients in each individual participant of the HCP (S-500) dataset, we assessed their functional meaning by testing whether they are related to recollection, a type of memory retrieval that the human hippocampus is known to be involved in. As predicted, we found that the TSM parameters summarising the overall spatial structure of the connectivity gradients predicted hippocampal-dependent recollection memory. Additionally, we tested whether the prediction of recollection memory required a gradient representation, or whether a characterisation in terms of parcels is sufficient (as suggested by Poppenk and Moscovitch, 2011). Our findings indicate that the gradient representation is more meaningful than a representation in terms of parcels when it comes to the prediction of recollection from the organisation of functional connectivity.

A 2014 review by Strange and colleagues already suggested that the dichotomous parcellation view, which has dominated the field for years, needs to be revisited as animal studies suggested that differences in connectivity between the hippocampus and other cortical and subcortical regions seem to be more gradual than abrupt. Although the most anterior and posterior parts of the hippocampus may have different functional specialisations, as suggested by studies linking behaviour to anatomical divisions (e.g., Poppenk & Moscovitch, 2011), there might not to be such a clear segregation of these parts. We found that, at least in the context of functional connectivity, a characterisation in terms of a gradient is more meaningfully related to recollection than a characterisation in terms of parcels. Our findings add weight to the idea that function is gradually distributed along the long axis of the hippocampus by providing the first in-vivo evidence from human neuroscience.

Our study is not the first to coin the idea that a gradient-like organisation might underlie the observed functional specialisation of anterior and posterior hippocampus. Recently, Persson and colleagues (2018) reported that episodic memory performance could be predicted from anterior, but not posterior resting-state functional connectivity, whereas the posterior resting-state functional connectivity was predictive of spatial memory. The authors explained these discrepancies by potential issues with disentangling the spatial and episodic components in their tasks, but they also pointed out another explanation, which emphasizes distribution of spatial representations along the entire long axis of the hippocampus, referring to a gradient of function. The evidence reported by Persson and colleages is based on the strength (not the organisation) of resting-state connectivity of a priori-defined seeds predicts hippocampal function, which is different from the question whether functional organisation within the hippocampus predicts hippocampal function. Nevertheless, both Persson and colleagues’ and our study point toward the idea that the characterisation of the functional organisation of the hippocampus in terms of a gradient is more meaningful than its characterisation in terms of parcels.

An open question is what mechanistic explanation underlies the result that the spatial organisation of the gradient estimated by resting-state functional connectivity predicts recollection. One possibility is that differences in the gradient reflect differences in the amount of neuronal resources that are dedicated to the task. Neurons that are dedicated to the same task likely exhibit similar connectivity fingerprints, and the gradient indicates which voxels exhibit similar connectivity fingerprints (i.e. similar colours in Figure 1). Thus, if recollection is poor in a participant, this participant might have fewer neurons with a particular connectivity fingerprint than a participant with good recollection, yielding different gradient maps.

More specifically, previous research has shown that connections from the neocortex to the hippocampus have preserved topographic organisation. The entorhinal cortex plays an important part as a relay in this process. It receives information from prefrontal cortex via topographically organized connections (Jones and Witter, 2007). We have previously demonstrated that within entorhinal cortex there is a topographic organisation that can be estimated using connectopic mapping (Navarro Schröder et al., 2015). This information in turn constitutes input to the hippocampus, which again exhibits topographic preservation. The implication is that differences in topographic organisation, as measured here, are indicative of differences in functional connectivity with the rest of cortex, potentially via topographic connections with entorhinal cortex. Although the present approach is limited to hippocampal-neocortical connectivity, it is likely that hippocampus displays similar gradients of connectivity with subcortical structures such as lateral septum (Risold and Swanson, 1996), amygdala (Kishi et al., 2006), and nucleus accumbens (Groenewegen et al., 1987). Future studies could elucidate whether individual differences in subcortical-hippocampal gradients predict motivated (e.g. reward-related) behaviours (Sheehan et al., 2004).

Unexpectedly, our results appear to suggest hemispheric differences, as the trend surface modelling parameters capturing the spatial organisation of the left connectivity gradient predicted face recollection, whereas the estimates of the right connectivity gradient (marginally) predicted recollection of places. The fact that our results did not show that recollection for faces can be predicted from the right hippocampus, and recollection for places by the left hippocampus does not necessary mean that these effects are not there, as our analyses might have been underpowered. However, it is also possible that our analysis is reflecting a true differentiation in hemispheric lateralisation. Previous studies have shown that damage to the right medial temporal regions, including the hippocampus, causes spatial memory impairments (Bohbot et al., 1998; Piggot & Milner, 1993), whereas similar damage in the left hemisphere affects primarily verbal memory (Bohbot et al., 1998; Milner, 1965). Though possible, these findings remain controversial, as other studies have shown that resections of either left or right temporal cortex produced impairments in spatial memory (Maguire et al., 1996). The observation that the topographic organisation subserving face recollection might be left lateralised resonates with the idea that face recollection might depend on concept forming, which in the broader context of face processing has been shown to be left-lateralised (Rangarajan et al., 2014). However, until these potential lateralisation effects are further scrutinized, these post-hoc accounts remain merely speculative.

In conclusion, we have shown that the functional organisation of the hippocampus is more appropriately described in terms of a functional gradient than in terms of functionally segregated parcels. We found that inter-individual differences in gradient are behaviourally relevant: the spatial organisation of the gradient along the hippocampal long axis (anterior-posterior) at an individual level predicted recollection, over and above the posterior-anterior ratio of the parcellation. Overall, these findings suggest that it is possible and meaningful to describe the functional organisation of the human hippocampus in terms of a gradient along its long axis. In addition, we have demonstrated that connectopic mapping approach is capable of mapping these gradients in individual subjects (albeit requiring high quality data; see Haak et al 2017), opening up the possibility to study how (aberrant) connectopic organisation of the hippocampus may underlie cognitive function in health and disease.

## Acknowledgements

This work was supported by the Netherlands Organization for Scientific Research Vidi Grant No. 864–12–003 (to CFB), Veni Grant No. 016.Veni.171.068 (to KVH), and Gravitation Programme Grant No. 024.001.006 (providing funds to IP). Data were provided by the Human Connectome Project, WU-Minn Consortium (Principal Investigators: David Van Essen and Kamil Ugurbil; 1U54MH091657) funded by the 16 NIH Institutes and Centers that support the NIH Blueprint for Neuroscience Research; and by the McDonnell Center for Systems Neuroscience at Washington University.

## Notes

**Conflict of Interest:** The authors declare no competing financial interests. CFB is Director and shareholder of SBGneuro Ltd.

